# FtsZ-mediated spatial-temporal control over septal cell wall synthesis

**DOI:** 10.1101/2024.01.29.577872

**Authors:** Longhua Hu, Amilcar Perez, Tanya Nesterova, Zhixin Lyu, Atsuhi Yahashiri, David S. Weisss, Jie Xiao, Jian Liu

## Abstract

FtsZ, the tubulin-like GTPase, is the central organizer of the bacterial divisome, a macromolecular complex that synthesizes new septal cell wall and degrades old septal cell wall (made of septal peptidoglycan, sPG) to allow cell wall constriction and cytokinesis. In *E. coli*, it is well accepted that 1) FtsZ recruits all essential divisome proteins to the septum, including the core sPG synthase complex, FtsWI/QLB and its activator, FtsN; 2) FtsWI/QLB must complex with FtsN to produce sPG under the wild-type background; and 3) the Brownian ratcheting by treadmilling FtsZ polymers drives the directional movements of sPG synthase proteins along the septum circumference; and 4) FtsZ is essential for the early stage, but dispensable for the late stage of cell wall constriction. However, it remains unclear how FtsZ spatial-temporally organizes the divisome for robust bacterial cytokinesis throughout cell wall constriction process. Combining theoretical modeling with experiments in *E. coli*, we show that at the early stage during cell division, the Brownian ratcheting by FtsZ treadmilling acts both as a template to corral FtsWI/QLB and FtsN into close contacts for FtsWI/QLB-FtsN complex formation and as a conveyor to maximally homologize the septal distribution of sPG synthesis activities to avoid uneven cell wall constriction. When the septum constricts progressively, the FtsN septal density increases via binding to denuded sPG; consequently, the denuded PG-bound FtsN serves as the template to activate FtsWI/QLB for continued sPG synthesis, rendering FtsZ dispensable. Our work establishes an overarching framework that FtsZ spatial-temporally controls over septal cell wall constriction.

**Significance:** Bacteria utilize FtsZ, the tubulin-like GTPase, to organize cell wall enzymes during cell division. FtsZ forms treadmilling polymers along the septum circumference and drives the directional movement of cell wall enzymes for robust cell wall constriction. How this role is achieved is unclear. We show that FtsZ treadmilling acts both as a template to corral cell wall enzymes into close contacts for priming and as a conveyor to homologize the septal distribution of cell wall synthesis activities for even septum constriction. These roles evolve at different stages of cell division and are modulated differentially by different bacteria; they likely define an overarching principle for robust cell division across the microbial world.

## Introduction

Cell division of most bacteria hinges on proper constriction of the septal cell wall, which entails the simultaneous synthesis of new septal peptidoglycan (sPG) and degradation of old cell wall circling the division plane (1, 2). FtsZ, the bacterial homolog of tubulin GTPase (3–5), is the central organizer of bacterial cytokinesis; it recruits more than a dozen of essential sPG enzymes and regulators (termed “divisome”) to the cell division plane (2, 6–8). However, it is not well understood how the activities of these sPG enzymes are spatially and temporally controlled to ensure proper cell wall constriction and robust cell division.

Temporally, the septal recruitment of divisome proteins follows a largely sequential order (2, 7, 9, 10). In *E. coli*, FtsZ, together with its membrane tethers and polymerization regulators FtsA and ZipA, first form a ring-like structure, termed Z-ring, at the future division site (11–13). The Z-ring then recruits key divisome components, including the core sPG synthase FtsWI/QLB complex (composed of five essential proteins, the sPG polymerase FtsW, the transpeptidase FtsI, and a tripartite subcomplex, FtsQLB) (2, 7, 9, 10, 14–17). In *E. coli*, new sPG synthesis does not commence until the arrival of an essential protein, FtsN, which also forms a ring-like structure termed the N-ring and activates the sPG synthases FtsWI by forming an FtsWI/QLB-FtsN complex (18, 19). As sPG synthesis proceeds, cell wall hydrolases (*i.e.*, amidases and lytic enzymes) hydrolyze old sPG to split the septum and shrink the septal diameter (*D*) (20–22). This process can be roughly divided into three stages based on the FtsZ and FtsN disassembly dynamics we and others previously reported (23, 24). In Stage I (1 μm > *D* > ∼ 0.7 μm), both FtsZ and FtsN assemble at the septum; the FtsZ level is significantly higher than that of FtsN. In Stage II (∼ 0.7 μm >*D* > ∼ 0.3 μm), the FtsZ-ring starts to disassemble while the FtsN-ring remains. In Stage III, (*D* < ∼ 0.3 μm), the FtsZ-ring is completely disassembled, and the FtsN-ring starts disassembling. Throughout the septum constriction process, FtsN recognizes and binds to denuded glycans, which are degradation products of sPGs generated by amidases (25–27). Because of its ability to activate new sPG synthesis and recognize degradation products, FtsN is believed to link sPG synthesis with hydrolysis (22, 28).

Spatially, FtsZ polymers treadmill at a speed of ∼ 30 nm/s in all bacterial species examined so far (29–33). In *E. coli*, treadmilling FtsZ polymers drive the directional movement of the essential sPG synthase complex (*e.g.*, FtsW and FtsI) along the septum perimeter (29–32) by a Brownian ratchet mechanism (34). Additionally, sPG synthase’s directional movements display a two-track feature (35): upon dissociating from the treadmilling FtsZ polymers (termed “Z-track”), FtsWI can stay stationary, diffuse away, or exhibit a slow, directional movement driven by processive new PG synthesis at a speed of ∼ 8-9 nm/s (termed “sPG-track”). FtsN has been shown to be critical to maintaining the processivity of sPG synthesis by forming a FtsWI/QLB-FtsN complex on the sPG track (28). FtsN associates with the Z-track at the beginning of cell wall constriction (Stage I); once the cell wall starts constriction (Stages II and III), FtsN becomes exclusively coupled to the sPG track or stays immobile off the tracks due to its binding to denuded glycan strands in the periplasm.

Many key questions remain unaddressed regarding the spatial-temporal control over sPG synthesis. For instance, past studies, including ours, show that the FtsWI and FtsQLB exist in a stable complex on both the Z and sPG tracks (16, 36). However, FtsWI/QLB and FtsN are expressed in low abundance (∼ 10s of molecules each) at septum (28, 37, 38), FtsWI/QLB join the divisome independent of FtsN (2, 7, 9, 10), and they spend most of their time on the Z-track and sPG track respectively without much free diffusion (if at all) (28, 35). The questions are: How do FtsWI/QLB and FtsN find each other and form the sPG synthase complex, FtsWI/QLB-FtsN, robustly and timely at septum? What, if any, is the role of FtsZ’s treadmilling dynamics in it? Further, when FtsZ polymers disassemble and become dispensable at a late stage (32, 39, 40), what maintains the spatiotemporal organization of sPG synthesis as cell division progresses? These unanswered questions lie at the heart of spatial-temporal control over septal cell wall remodeling.

To address these questions and shed light on the organizational principles of bacterial cell division, we combined theoretical modeling with experimental testing using *E. coli* as the model system. We found that FtsZ treadmilling promotes the FtsWI/QLB-FtsN complex formation by “corralling” the molecules into close and frequent contacts, providing an advantage over the diffusion-and-collision mechanism. At the beginning of cell wall constriction (Stage I), the FtsZ treadmilling speed is optimal for evenly distributing formed FtsWI/QLB-FtsN complexes and, hence, the sPG synthesis activities along the septum circumference. As the septum constriction progresses (Stages II and III), FtsN molecules concentrate at the septum due to their sustained binding to denuded PGs. The higher level of FtsN then increases the frequency of FtsWI/QLB-FtsN complex formation, which not only sustains sPG synthesis and speeds up the septal constriction rate but also prevents FtsWI/QLB complexes from being released to the Z-track - they are either retained on the sPG track to synthesize sPG or remain bound to denuded glycan-anchored FtsN molecules. Thus, in the late stages, FtsN renders FtsZ dispensable. As such, while FtsZ treadmilling facilitates the formation of FtsWI/QLB-FtsN complex and smoothens out the sPG synthesis activity along the septum, FtsN takes the baton from FtsZ as the septum constriction progresses toward completion. Together, our work delineates the evolving roles of FtsZ in controlling the spatial-temporal distribution of sPG synthesis as cell division progresses, paving the way to elucidate the organizational principles for robust bacterial cell wall constriction.

## Results

### FtsZ treadmilling increases the frequency of close contacts between FtsWI/QLB and FtsN

To investigate how FtsWI/QLB and FtsN may robustly form the core divisome complex at the septum despite their low copy numbers at an early stage (Stage I) of cell wall constriction, we first leveraged a previous agent-based theoretical modeling in *E. coli* to examine the likelihood that FtsWI/QLB and FtsN are in close contact, the prerequisite for FtsWI/QLB-FtsN complex formation. We explore two scenarios (**Fig. 1A**): In the first scenario (left), there is no FtsZ, and FtsWI/QLB and FtsN randomly diffuse and collide. In the second scenario (right), FtsZ treadmilling drives directional movements of FtsWI/QLB and FtsN via a Brownian ratchet mechanism (34).

**Figure 1.**
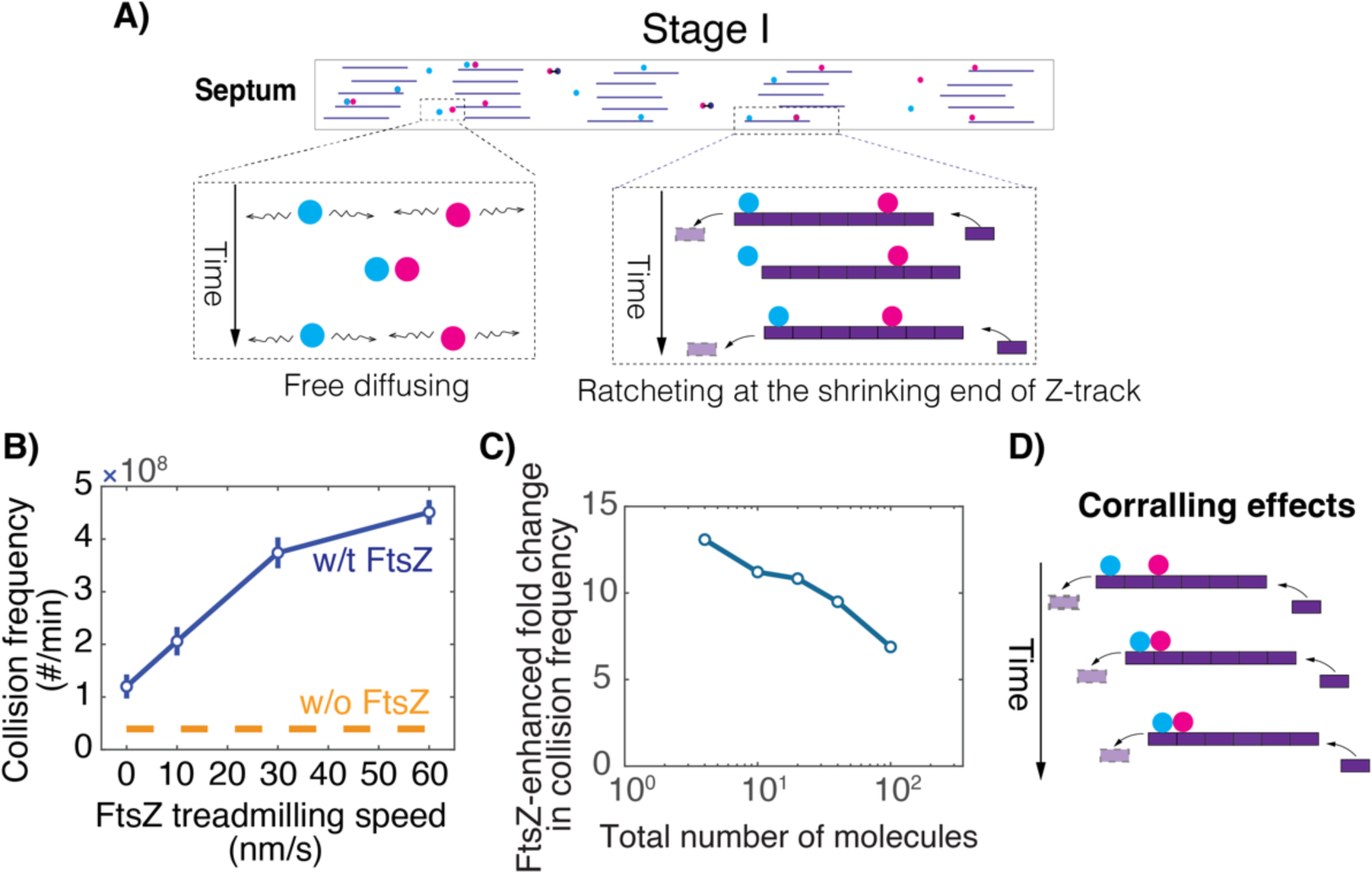
Treadmilling of FtsZ polymers promotes the collision frequency of FtsWI/QLB and FtsN. A) Schematic of model setup describing the interactions between treadmilling FtsZ polymers (purple), FtsWI/QLB (blue), and FtsN (pink). B) FtsZ treadmilling promotes the collision frequency of FtsWI/QLB–FtsN close contacts. C) The enhancement in the collision frequency by FtsZ treadmilling decreases with the number of FtsWI/QLB and FtsN molecules. Hereby, this enhancement is defined by the ratio of the collision frequency per min between the cases with and without FtsZ; for the case with FtsZ, the treadmilling speed is fixed at 30 nm/s. The number of molecules refers to the sum of FtsWI/QLB and FtsN molecules with 1:1 ratio. D) Schematic describing FtsZ treadmilling-mediated corralling effects.

The model in this section focuses on Stage I of septum constriction. It depicts the septum as a 2D domain of 3000 nm long and 100 nm wide based on previous superresolution imaging studies (23, 41–43). It imposes periodic and hard-wall boundary conditions lengthwise and widthwise, respectively. 20 FtsWI/QLB molecules and 20 FtsN molecules are randomly distributed within the septum. In the later simulation for Stages II and III of septum constriction, FtsN molecule numbers will evolve between 20 and 60 as septum constriction progresses (28), whereas FtsWI/QLB complex numbers will be at 20. For simplicity, the model treats FtsWI/QLB as one complex, as previous studies, including ours, showed that FtsWI and FtsQLB form a stable complex without other divisome components (16, 36). Each FtsWI/QLB complex or FtsN molecule is represented by a single circular particle of 5 nm in diameter (44–46), and they volumetrically exclude each other.

In the first scenario (**Fig. 1A**, left), there are no FtsZ polymers in the septum. FtsWI/QLB and FtsN diffuse freely as membrane protein complexes (∼ 0.04 μm^2^/s) (28, 34) in the region until they collide. In the second scenario (**Fig. 1A**, right), FtsWI/QLB and FtsN bind to FtsZ via the short-ranged binding potentials; they will either end-track the shrinking end via the Brownian ratchet mechanism (34), or be stuck in the middle of treadmilling FtsZ polymers (**Movie S1**). When these particles dissociate from FtsZ polymers, they diffuse freely as membrane protein complexes (∼ 0.04 μm^2^/s) (28, 34), until they collide with each other or interact with FtsZ again. For this scenario, we model five bundles of FtsZ polymers treadmilling from left to right at 30 nm/s along the septum; each bundle consists of five FtsZ polymers 200 nm long, with each FtsZ subunit modeled as a circular disk of 5 nm in diameter. Because both FtsWI/QLB and FtsN bind to FtsZ through interactions mediated by FtsA-binding potential (47–50), the FtsZ-binding potential is modeled as the same harmonic potential for FtsWI/QLB and FtsN with the potential depth of ∼ 10-20 *k*_B_T (corresponding to a dissociation constant *K_d_* in nM to μM range) and the short range of ∼ 3-10 nm, commensurate with size of FtsZ subunit, similar to what we have described previously (51–54).

We next leveraged stochastic simulations of this agent-based model to determine how FtsZ treadmilling modulates the likelihood that FtsWI/QLB and FtsN are in close contact: 1) the first-passage-time (FPT), which is the time it takes for one FtsWI/QLB to collide with an FtsN (or *vice versa*) for the first time; and 2) the collision frequency, which is the number of times FtsWI/QLB and FtsN collide with each other per unit time. Here, we defined a collision event as the center-to-center distance between FtsWI/QLB and FtsN being less than a threshold distance, the nominal value of which was chosen to be 10 nm. For the model results presented below, the essential conclusion remains robust against parameter variations such as this threshold distance (**Fig. S1A**) and protein diffusion constants (**Fig. S1B**).

Our model results suggest that while the FPT remains fast regardless (in the millisecond range) (**Fig. S2**), FtsZ’s treadmilling significantly enhances the collision frequency between FtsWI/QLB and FtsN as compared to the free-diffusing case. The enhancement increases with the FtsZ treadmilling speed (**Fig. 1B**). Notably, even in the regime where FtsZ does not treadmilling, *i.e.*, the speed is 0 nm/s, we still observed an enhancement of FtsWI/QLB-FtsN complex formation. This phenomenon is because FtsZ polymers can still act as a landing pad to enrich the binding of FtsWI/QLB and FtsN while allowing their 1-D diffusion along the FtsZ polymers, which increases the collision frequencies compared to the free diffusion scenario. We further showed that, as the number of FtsWI/QLB and FtsN increases from 4 to 100 (with 1:1 ratio), the enhancement decreases from ∼ 14 folds to ∼ 6 folds at the treadmilling speed of 30 nm/s (**Fig. 1C**). Given that the septal levels of FtsWI/QLB and FtsN are typically < 100 in *E coli.* (28, 37, 38, 55) (also see the parameter determinations in Table S1), our findings suggest that FtsZ treadmilling potentiates the formation of FtsWI/QLB-FtsN complex by increasing the collision frequency. These results point to the following physical picture (**Fig. 1D** and **Movie S2**): As an end-tracking FtsWI/QLB particle persistently approaches an FtsN molecule stuck in the middle of the same treadmilling FtsZ polymer (or *vice versa*), FtsZ’s treadmilling essentially corrals them into close contact with each other. Since FtsZ-binding affinities provide a template effect preventing both molecules from diffusing too far away, FtsWI/QLB and FtsN will be in close contact more frequently than the free-diffusing case. The more frequent close contacts between FtsWI/QLB and FtsN, the faster the complex will form. Therefore, our model results suggest that the FtsZ treadmilling facilitates the formation of FtsWI/QLB-FtsN complexes.

### Two-track model of sPG synthesis

Next, to describe how FtsZ’s treadmilling dynamics may control the spatial-temporal activities of sPG synthases, we expanded the above theoretical modeling to dissect the two-track feature of sPG synthesis (**Fig. 2A**) as recently observed in *E. coli* (35). The model integrates three essential steps of sPG synthesis. In the first step, when FtsWI/QLB and FtsN are in close contact (as defined in **Fig. 1**), they will reversibly form an intermediate complex, FtsWI/QLB-FtsN^‡^ with on and off rate constants (*k*_on_, *k*_off_), like any biochemical binding reaction. FtsWI/QLB-FtsN^‡^ will then be converted to the primed, stationary sPG synthase complex, FtsWI/QLB-FtsN*, with a priming rate of *k*_prime_ (**Fig. 2A**), *i.e.*,

**Figure 2.**
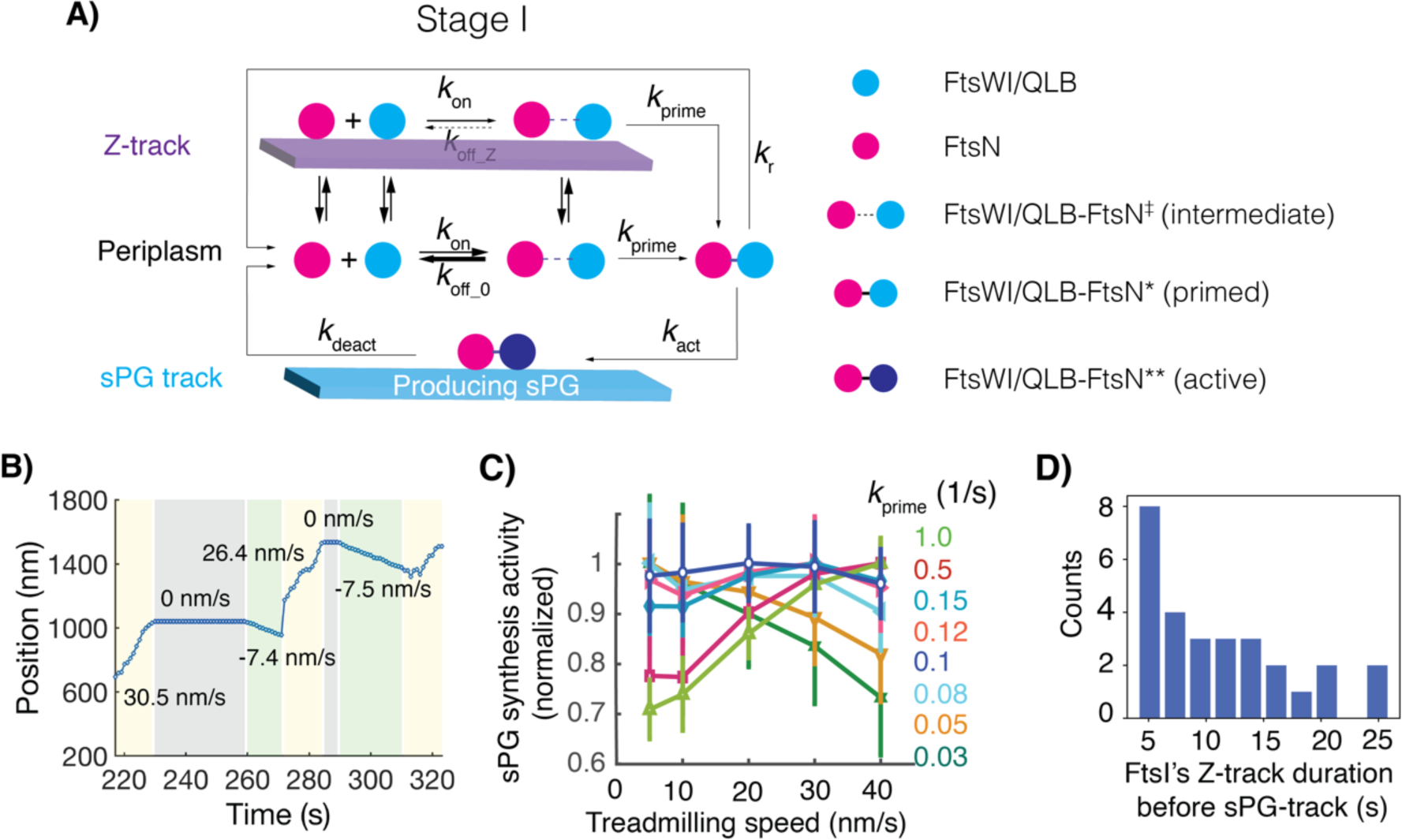
FtsZ treadmilling spatial-temporally controls sPG synthesis activity in two-track model. A) Schematic of two-track model that consists of three sets of reactions. The first set of reactions is priming sPG synthase: FtsWI/QLB and FtsN reversibly form an intermediate complex, FtsWI/QLB–FtsN^‡^ with on and off rates, *k*_on_ and *k*_off_. When this reaction is on Z-track, the off rate *k*_off_Z_ is typically chosen to zero; otherwise, the off rate, *k*_off_0_ is very fast (typically ∼ 10^3^/s in the model). The FtsWI/QLB–FtsN^‡^ complex will then convert the primed sPG synthase, FtsWI/QLB– FtsN* with the priming rate of *k*_prime_. The second set of reactions is forming stationary FtsWI/QLB-FtsN* complex: Once formed, the FtsWI/QLB-FtsN* will immediately dissociate from the Z-track and remain stationary probably by binding to denuded PG. While this FtsWI/QLB-FtsN* waits for the recruitment of lipid II, the building block of cell wall, it has two fates of either reverting to free-diffusing FtsWI/QLB and FtsN at the rate of *k*_r_∼ 0.1/s or being activated into FtsWI/QLB-FtsN** at the rate of *k*_act_ ∼ 0.1/s for sPG synthesis. The latter represents the kinetics that lipid II recruitment to the FtsWI/QLB-FtsN*. The third set of reactions is sPG synthesis: The activated FtsWI/QLB-FtsN** will synthesize sPG in either the left or right directions with 50% of probability each. However, once initiated in one direction, the sPG synthesis will persist in that direction at the speed of ∼ 8 nm/s with the duration of ∼ 10 – 20 seconds, beyond which the enzyme will fall apart into the free-diffusing FtsWI/QLB and FtsN in periplasm. The activated sPG synthase complex (FtsWI/QLB-FtsN**) is assumed to move directionally at the same speed of sPG synthesis. B) A representative simulation trajectory of FtsWI/QLB in two-track model. The speed of FtsWI/QLB is marked in each segment of the trajectory. C) Predicted dependence of sPG synthesis rate on FtsZ treadmilling speeds with different *k*_prime_ rates. D) Single-molecule tracking experiments reveal a slow formation of FtsWI/QLB-FtsN* on Z-track. This histogram quantifies the lifetime of the FtsW that end-tracks FtsZ polymers (the fast track) right before generating sPG (the slow track).

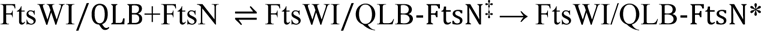

The primed state is supported by our single-molecule tracking observation that the transition of a FtsW molecule between the Z-track and sPG-track often goes through an immobile state with a characteristic lifetime (35), which likely reflects a primed state of the synthase complex poised for a signal for activation or inactivation in the periplasm. Here, the on-rate constant *k*_on_ depicts the intrinsic rate of complex formation, regardless of whether the reaction is on the Z-track or in the periplasm. Due to the enhanced collision frequency by the treadmilling FtsZ polymers (**Fig. 1B**), the level of the intermediate complex formation, FtsWI/QLB-FtsN^‡^, is expected to increase on Z-track significantly. The off-rate constant *k*_off_, however, is expected to be modulated by the Z-track because of previously reported interactions of FtsW, FtsI, and FtsN with FtsZ, which are mediated through FtsA and ZipA (11, 47–50). If FtsWI/QLB-FtsN^‡^ is FtsZ-bound, the off-rate constant (denoted as *k*_off_Z_) is small, and the complex is stable; if not, the FtsWI/QLB-FtsN^‡^ will revert to FtsWI/QLB and FtsN rapidly with a large off rate constant (denoted as *k*_off_0_). While precise values of *k*_on_, *k*_off_0_, *k*_off_Z_, and *k*_prime_ are unknown, we calculated phase diagrams of these parameters and identified the parameter regime that satisfied the observed insensitivity of total sPG synthesis activity to FtsZ’s treadmilling speed (29) (**Fig. S3**). In the text below, if not otherwise mentioned, the choice of model parameters will be all within this nominal parameter regime so that the model results are constrained by our experiments.

In the second step, once formed, the primed complex FtsWI/QLB-FtsN* will immediately dissociate from the Z-track into the periplasm because of the binding of FtsN’s SPOR domain to denuded glycans (25–27), where the complex likely remains stationary, in accordance with our single-molecule tracking studies (35). For simplicity, the model does not explicitly describe the denuded glycan-binding of FtsN. A primed FtsWI/QLB-FtsN* complex has two independent fates: it can either initiate sPG synthesis (denoted as the active complex FtsWI/QLB-FtsN**) upon the availability of lipid II, the cell wall precursor, at a rate of *k*_act_, or revert to free-diffusing FtsWI/QLB and FtsN particles at a rate of *k*_r_. Based on our single-molecule tracking studies (28, 35), we assign *k*_act_ and *k*_r_ both at ∼ 0.1s^-1^.

In the last step, once sPG synthesis initiates, a FtsWI/QLB-FtsN** complex will run processively, with equal probability of moving in the forward or backward directions. Once a sPG synthesis event starts, however, it will persist in one direction for on average ∼ 20 s at the speed of ∼ 8 nm/s, based on our single molecule tracking measurements (28, 35). When the sPG synthesis stops, the enzyme will fall apart into free-diffusing FtsWI/QLB and FtsN particles, which will either interact with FtsZ and/or each other, initiating another round of events as described in the three steps. If not otherwise mentioned, we calculate the total sPG synthesis activity in the septum region using the total sPG length synthesized over 15 min: *i.e.*, the sum of all sPG track lengths, which is equal to the speed of sPG track (8 nm/s) times its duration.

With the three steps, the model depicts the chemical interconversions between different states of sPG synthases, which are modulated by their interaction kinetics with each other, FtsZ, and sPG synthesis activity. We next used stochastic simulations to reveal the dynamic nature of these spatial-temporal controls, with most of the kinetic parameters measured by our experiments (ref, see the Model Parameter Table S1 in SI). **Fig. 2B** shows one representative simulation trajectory of FtsWI/QLB-FtsN in the two-track model. One sPG synthase FtsWI/QLB alternates between the fast-moving Z-track and the slow-moving sPG track, interspersed by two stationary states with one stuck in the middle of treadmilling FtsZ polymers and the other binding to FtsN waiting for the full activation. Due to the high density of FtsZ and the high binding potential, the enzymes display minimal diffusive motions. These behaviors recapitulate the observed tracking and transitioning dynamics of FtsWI/QLB on the two tracks in our experiments (28, 35).

### FtsZ treadmilling speed can have opposing roles in controlling total sPG synthesis activity

Cell wall constriction speed was reported to be sensitive to FtsZ treadmilling speed in *B. subtilis* (30) but not *E. coli* or *S. pneumoniae* (29, 31). Our previous 1-D Brownian ratcheting model suggests that a faster treadmilling speed of FtsZ could lead to a higher fraction of cell wall enzymes exiting the Z-track to become available for sPG synthesis (34). The extent of this effect may depend on the differential binding potentials of cell wall enzymes to the Z-track in these different bacterial species (34). Hereby, our two-track model adds new insights into these observations. The corralling effect of treadmilling FtsZ polymers allows for a more frequent FtsWI/QLB-FtsN^‡^ intermediate complex formation at a faster treadmilling speed and hence potentially more primed complex FtsWI/QLB-FtsN*, higher sPG synthesis activity, and faster cell wall constriction speed. As such, depending on the value of the priming rate constant *k*_prime,_ the FtsZ treadmilling speed could affect the total sPG synthesis activity differentially. Indeed, as shown in **Fig. 3C**, we found that when *k*_prime_ is large (> 0.15 s^-1^), sPG synthesis increases almost linearly with FtsZ treadmilling speed. In contrast, when *k*_prime_ is small (< 0.05 s^-1^), sPG synthesis decreases with the FtsZ treadmilling speed. Only with an intermediate value (0.05 s^-1^ < *k*_prime_ < 0.15 s^-1^), does the sPG synthesis rate remain insensitive to the FtsZ treadmilling speeds between 5 and 30 nm/s, akin to the observation in *E. coli* (29). These predictions remain robust against varying model parameters (**Fig. S3**). Notably, given that most sPG synthases are primed *via* Z-track before they can produce sPG based on the model, the dwell time of sPG synthases on the fast Z-track right before they switch to the slow sPG track or become immobile could be used as the proxy to infer the reaction time. Using this criterion, we quantified FtsI’s average dwell time on the Z-track before it switches to the slow sPG track or becomes immobile at 12 ± 6 s based on our single-molecule tracking data (**Fig. 2D**). Therefore, we inferred that the priming rate of sPG synthase is ∼ 0.06-0.16/s, supporting the model prediction.

**Figure 3.**
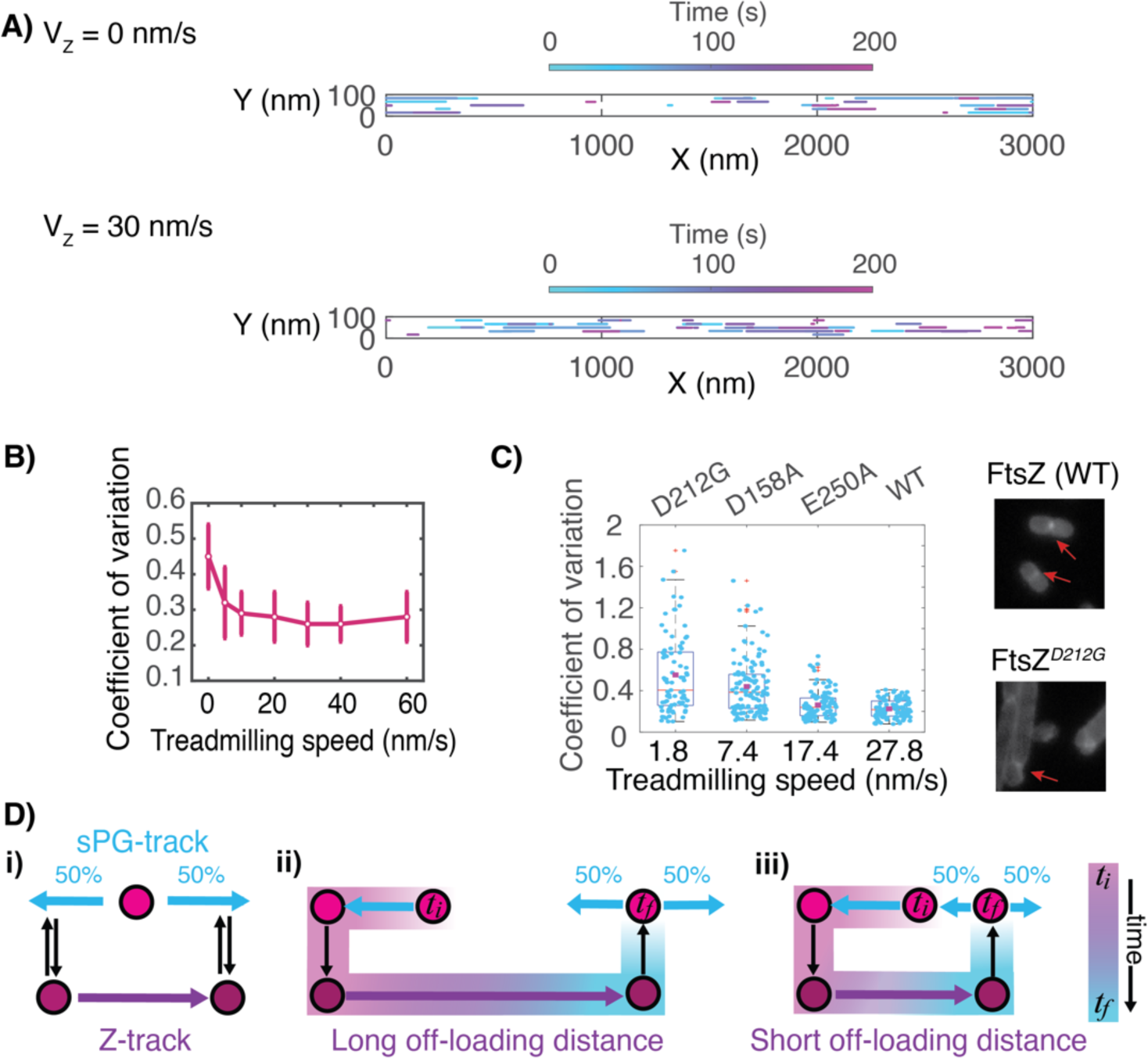
Predicted spatial distribution of sPG synthesis as a function of FtsZ treadmilling speed. A) Representative spatial profiles of newly synthesized sPG strands in the septum at different FtsZ treadmilling speeds. *Above*: 0 nm/s. *Below*: 30 nm/s. The color of the trajectory denotes the timing of the sPG strand synthesis. For clarity of presentation, only one tenth of the trajectories is plotted for each case. B) Predicted dependence of spatial heterogeneity in sPG synthesis on FtsZ treadmilling speeds. C) HADA labeling reveals the spatial distribution of newly synthesized sPG with different FtsZ GTPase mutants in *E. coli* that slow down the FtsZ treadmilling speed, as indicated on the x-axis. D) Schematic describing the role of FtsZ treadmilling-mediated positioning in promoting even distribution of sPG synthesis activities along the septum. Here, a sPG synthase complex is represented by a circular disk, which transitions between different states on Z-track and sPG track. The thick cyan arrow marks both the potential direction and the length of the sPG synthesis in which the left and the right directions have the equal probability of 50%, which is schematized in (i). Comparisons between (ii) and (iii) illustrates that the newly synthesized sPG strands tend to cluster together when the off-loading distance (marked by the thick magenta arrows) is shorter, and hence distribute more heterogeneously.

Taken together, we suggest FtsZ’s treadmilling speed has opposing roles in controlling sPG synthesis activity. Due to the corralling effects, the faster the FtsZ treadmills, the more frequently the FtsWI/QLB and FtsN will contact each other to form the intermediate FtsWI/QLB-FtsN^‡^ complex (**Fig. 1B**). However, the faster FtsZ treadmills, the less time FtsWI/QLB and FtsN will remain on the Z-track as we previously showed both experimentally and theoretically (34), and hence will be less likely in close contact to form the primed FtsWI/QLB-FtsN* complex. Therefore, the frequency and the duration that FtsWI/QLB and FtsN are in close contact hinge on FtsZ treadmilling speeds oppositely; only at the intermediate priming rate constant range (0.05-0.15 s^-1^), these two opposite effects balance out, and the sPG synthesis rate remains insensitive to FtsZ treadmilling speed.

### FtsZ treadmilling speed is optimized to distribute sPG synthesis evenly along the septum

A robust cell wall constriction process entails evenly distributed sPG synthesis along the septum so that the septum can constrict concentrically (29). Previously, we observed that FtsZ’s treadmilling dynamics dictate the spatial distribution of sPG synthesis activities along the septum. But how this effect is achieved is unclear. Using the two-track model, we examined the spatial profile of sPG synthesis activity (**Movie S3**) and how it depends on FtsZ treadmilling speed.

**Fig. 3A** presents two representative spatial profiles of newly synthesized sPG strands along the septum over time: one without FtsZ treadmilling (speed at 0 nm/s) and the other at 30 nm/s, with all other parameters kept the same. The spatial distribution of sPG synthesis activity without FtsZ treadmilling is more heterogeneous along the septum than with the FtsZ treadmilling speed at 30 nm/s. To quantify how uniformly the sPG synthesis distributes, we calculated the coefficient of variance (CV) of the density of newly synthesized sPG strands along the septum. The higher the CV, the more unevenly sPG synthesis activity is distributed. The model predicts that the CV is sharply dependent on FtsZ’s treadmilling speed when it is slower than 20 nm/s while remaining low (*aka* uniformly distributed) at the wild-type FtsZ treadmilling speed of ∼ 20 – 60 nm/s (**Fig. 3B**).

To verify this prediction, we used our previous experimental measurements where the spatial distribution of newly synthesized sPG along the septa was visualized by an 8-min pulse labeling of HADA, a fluorescent D-Ala-D-Ala analog, in a set of *E. coli* FtsZ GTPase mutants that have different treadmilling speeds (29). The calculated CV shows that with decreasing FtsZ treadmilling speeds (from ∼30 nm/s to ∼ 2 nm/s), the distribution of newly synthesized sPG intensity along the septum became increasingly heterogeneous (**Fig. 3C**), despite that their total amount remains relatively constant as we previously reported (29). Consequently, instead of concentric constriction, the cell septum constricted asymmetrically, and the cell division was compromised. These findings support the notion that FtsZ treadmilling speed may be optimized for robust cell wall constriction in *E. coli* by evenly distributing new sPG synthesis activities and hence allowing septum constriction concentrically, consistent with our previous conclusion (29).

Taken together, we suggest the following picture of FtsZ-mediated spatial-temporal controls over sPG synthesis (**Fig. 3D**): Because FtsWI/QLB-FtsN* can only synthesize sPG when it is off the Z-track, it can be considered that a treadmilling FtsZ polymer acts as a conveyor to keep a FtsWI/QLB-FtsN^‡^ intermediate complex inactive while transporting it until the intermediate complex becomes the primed FtsWI/QLB-FtsN* complex, which then exits the Z-track (**Fig. 3Di**). Therefore, the persistent run length of sPG synthase enzymes on the Z-track is the off-loading distance that gauges how far the complex is kept inactive along the septum; it decreases as the FtsZ treadmilling speed decreases from 30 nm/s, as demonstrated in our previous modeling work (34). Our current model shows that the longer the off-loading distance is, the more evenly the activation sites of a sPG synthase complex and hence the newly synthesized sPG strands spread out along the septum (the cyan arrow in **Fig. 3Dii**). Hence, the FtsZ treadmilling speed in wild-type *E. coli* maximizes the off-loading distance and helps deposit the newly synthesized sPG along the septum evenly. Conversely, decreasing the FtsZ treadmilling speed in *E. coli* shortens the off-loading distance and increases the probability of unevenly distributed new sPG synthesis sites along the septum (**Fig. 3Diii**, also see discussion).

### FtsZ is dispensable in sPG synthesis at the late stage of cell division

Recent experiments in *E. coli, S. aureus,* and *B. subtilis* revealed that while essential for the initial assembly of the divisome, FtsZ polymers start to disassemble midway and become dispensable for cell wall constriction at late constriction stages (32, 39, 40). How does the decrease in the FtsZ level during the late stages of septum constriction impact the spatial-temporal regulations of sPG synthesis? What guides the continual and smooth sPG synthesis to complete the cell wall constriction without FtsZ or its treadmilling activity? These questions lie at the heart of FtsZ-mediated divisome regulation and present a perfect testing ground for our model. If it holds up, the model should be able to explain these disparate roles of FtsZ in cell division coherently and quantitatively.

We quantitatively adapted our model to the experimentally measured FtsZ and FtsN septal intensity as a function of the septum dimension (23, 24) (**Fig. 4A**) during stages I, II, and III of cell wall constriction. We used our previous measurements of cell wall constriction speed to simulate the decrease of the septum diameter over time (23). The model depicts that in Stage I (0.7 < *D* < 1 μm), there are 25 FtsZ polymers of 200 nm-long and 20 FtsN molecules with no denuded sPG, which is the model setup in **Figs. 1-3**. In Stage II (0.3 < *D* < 0.7 μm), the number of FtsZ polymers gradually decreases from 25 to 10 (41), during which there are 60 FtsN molecules (28) that are either bound to denuded sPG and hence remaining stationary, or engaged in the FtsWI/QLB-FtsN** complex processively synthesizing sPG without binding to denuded glycans. In Stage III, (*D* < 0.3 μm), 30 FtsN molecules remain (28) with no FtsZ (23).

**Figure 4.**
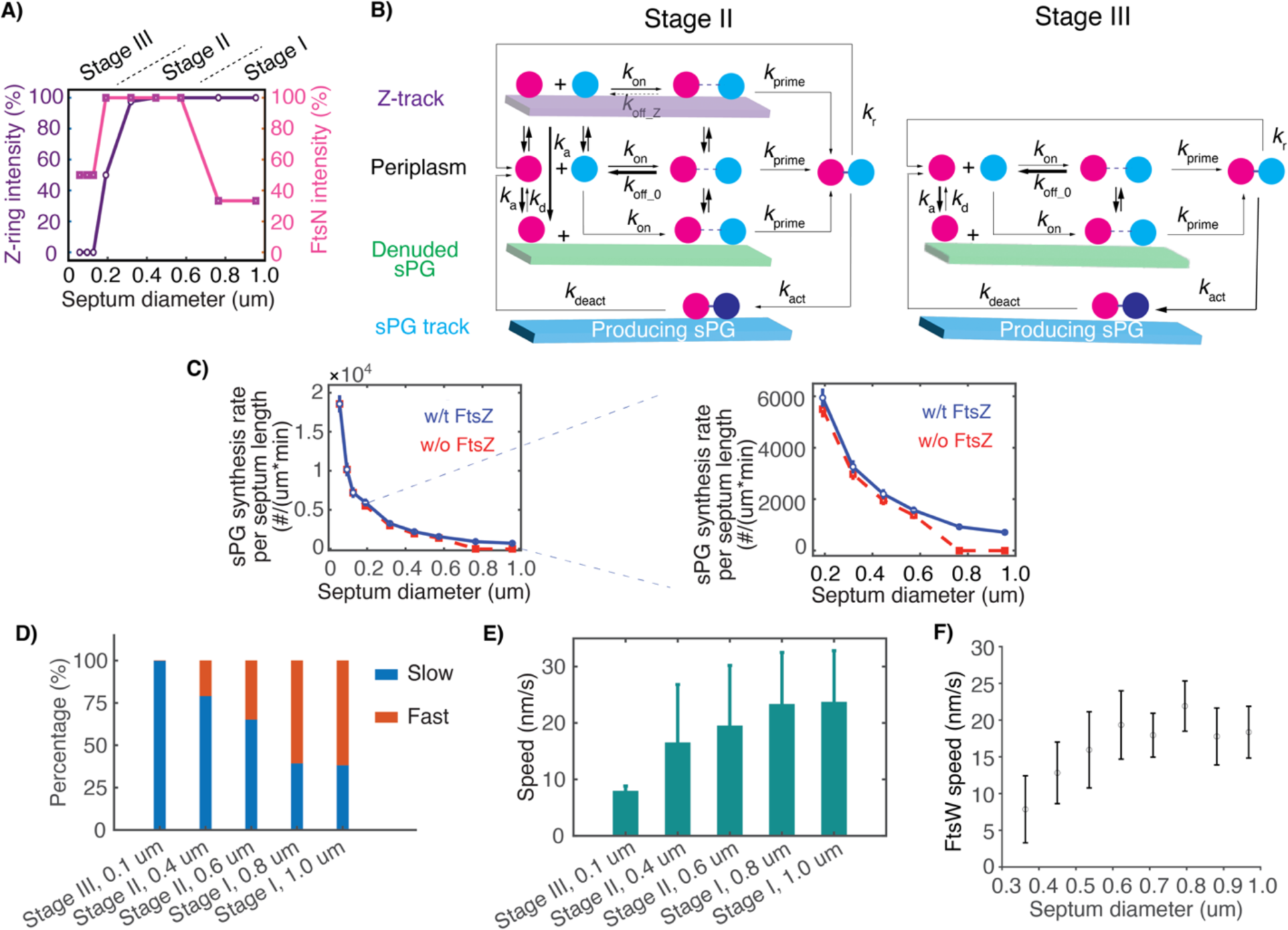
FtsN takes the baton from FtsZ in promoting sPG synthesis as septum constricts. A) Measured changes in FtsZ and FtsN septal levels as septum constricts in *E. coli*. B) Schematics of model depicting the divisome dynamics at the stages II and III of cell division. C) sPG synthesis rate per septum length is predicted to increases as a function of septum diameter. The blue and red curves correspond to the cases with and without the presence of FtsZ, respectively. Note that, the case with the presence of FtsZ mimics the WT case, in which the evolution of FtsZ septal level follows that in Fig. 4A and the corresponding treadmilling speed is kept at 30 nm/s. The case without the presence of FtsZ refers to the scenario that does not have FtsZ nor the treadmilling throughout the Stage I, II and III of septum constriction. *Left*: The entire process of septum constriction (Stage I to Stage III). *Right*: The zoom-in view of Stage I and Stage II. D) The model calculation shows that the sPG synthase complex spends more and more time on sPG track as the septum constricts. E) The average speed of sPG synthase’s directional movement as the function of the septum diameter. F) SMT measurement of the FtsW’s directional speed at different septum diameters in *E. coli*.

While keeping the parameters of all the chemical reactions the same as those in Stage I, the model in Stages II and III additionally incorporates the following experimental observations (**Fig. 4B**). First, in Stages II and III, an FtsN molecule will bind to denuded glycans at a rate constant of *k*_a_, significantly higher than that resulting from the binding potential between FtsN and FtsZ, in accordance with our single-molecule tracking experiments, which showed that no FtsN moves on the Z-track during these stages. The essence of the model result presented below holds as long as the FtsN-denuded PG binding is sufficiently tight (**Fig. S4**), in line with our experimental measurements using purified sacculi (**Fig. S5**) and previous report (25–27). Second, based on our single-molecule tracking data, FtsN molecules bound to denuded glycans are stationary and have an average lifetime of ∼ 27 s (28). Third, once a diffusing FtsWI/QLB complex in the periplasm engages with a denuded sPG-bound FtsN, the resulting FtsWI/QLB-FtsN^‡^ intermediate complex will persist until it converts to the primed complex, FtsWI/QLB-FtsN*.

Integrating the stochastic model simulation results from Stage I to III provides the first coherent explanation for the observed diminishing role of FtsZ as septum constriction progresses (**Figs. 4C-F**): 1) the total sPG synthesis activity per unit area increases as the septum constricts, in accordance with our previous observation that the cell wall constriction speed accelerates (23); 2) FtsZ is only essential for even sPG synthesis distribution in Stage I (32, 39, 40) while depleting FtsZ from the system in the later stages of cell division does not alter the septum constriction speed notably (**Fig. 4C**). The different requirement for FtsZ is because in Stages II and III, as the septum constricts, it concentrates FtsWI/QLB and the even more FtsN molecules (*via* binding to denuded PG), both of which increase collision frequency between FtsWI/QLB and FtsN. The increased collision enhances the FtsWI/QLB-FtsN* complex formation, sustains sPG synthesis activity, speeds up the septal constriction rate, and renders FtsZ dispensable. Consequently, the model predicates that during late stages of septum constriction, sPG synthases would spend most of their time binding to the denuded PG-bound FtsN or producing sPG instead of on Z-track (**Fig. 4D**), leading to a decreased average speed of sPG synthases’ directional movements as the cell division progresses (**Fig. 4E**). To test this prediction, we used our previous SMT experiments to quantify how the partitioning of fast and slow-moving populations of FtsW evolve as the cell division proceeds. Indeed, we observed that as the septum constricts majority of FtsW molecules moved slowly, leading to a decreased average speed of directional movement (**Fig. 4F**). As such, our work establishes a coherent model that describes the evolving roles of FtsZ in sPG synthesis and cell division: as the septum constriction progresses, the FtsZ treadmilling-mediated corralling effect pass the baton to the denuded PG-bound FtsN in activating sPG synthases.

## Discussion

In this study, we presented evidence that treadmilling FtsZ polymers not only promote the formation of the sPG synthase complex by corralling its essential components into frequent close contacts but also control the positioning of active sPG synthase complexes along the septum. Specifically, at the early stage of cell division, the presence of FtsZ is essential for activated sPG synthase complex formation; however, the total amount of synthesized sPG is insensitive to FtsZ treadmilling speed. Rather, the FtsZ treadmilling speed in wild-type *E. coli* maximizes the off-loading distance of sPG synthase and evenly distributes the sPG synthesis activities along the septum, underlying a robust and concentric septal constriction. At the late stage of cell division, however, the FtsZ-ring disassembles and becomes nonessential for septal constriction regarding the spatial distribution and the total sPG synthesis activity. Instead, the denuded PG from the hydrolysis of old sPG potentiates the septal accumulation of FtsN that takes the baton from FtsZ to activate sPG synthase and complete the septum constriction. Our finding establishes an overarching framework of septal cell wall synthesis, elucidates the evolving roles of treadmilling FtsZ polymers in regulating sPG synthesis as the septum constricts, dissects the coordinated actions between FtsZ and FtsN for sustained septal constriction, and paves the way toward the coherent mechanism of spatial-temporal controls over the divisome dynamics and, hence, cell division.

While interrogating the spatial-temporal dynamics of the entire divisome will require future work, our results begin to shed new light on the molecular logic behind the inner workings of the divisome. For instance, *E. coli* cells have a single layer-cell wall and, therefore, need to control the timing and location of sPG synthesis tightly, in synchrony with genome partition and other cell-cycle events. To achieve this goal, *E. coli* seems to adopt multiple regulations to prevent premature activation of sPG synthesis. First, FtsWI by itself only exhibits minimal sPG synthesis activity (14). Second, FtsQLB is required to scaffold the FtsWI complex (16), which will gain full activation upon binding with FtsN (18, 19). Third, the amounts of FtsWI/QLB and FtsN complexes are low at the septum (28, 37). Here, we delineated two additional regulatory layers: 1) To cope with the scarcity of FtsWI/QLB and FtsN molecules for a timely sPG synthesis, treadmilling of FtsZ polymers not only potentiates the formation of primed FtsWI/QLB-FtsN* complex but keeps the resulting sPG synthase complex inactive; 2) only the FtsWI/QLB-FtsN* complex that dissociates from Z-track will be able to produce sPG (35). In this Z track-mediated regulation, a key rate-limiting step of sPG synthesis is the slow conversion of intermediate complex FtsWI/QLB-FtsN^‡^ to the primed complex FtsWI/QLB-FtsN*, which takes on average ∼ 12 s on the Z-track in *E. coli* (**Fig. 2**). *E. coli* cells exploit this slow conversion process to allow even deposition of the thin layer of newly synthesized sPG along the septum (**Fig. 3**), which is necessary to maintain cell wall integrity (56).

Different bacteria may exploit the FtsZ-mediated regulation of sPG synthesis distinctively to cope with their own functional needs. For instance, if the formation of primed sPG synthase complex is fast, then the faster the FtsZ polymers treadmills, the faster the sPG synthase will dissociate from the Z-track and, hence, the faster will be the sPG synthesis (**Figs. 2C & S3D**). This scenario may account for the cell division of a bacteria with a thick cell wall (*e.g.*, *B. subtilis*), which, unlike *E. coli*, may favor a fast sPG synthesis rate over the precise spatial distribution of sPG deposition, since the sPG in *B. subtilis* is disordered as demonstrated by AFM studies (57). According to our model, the observed increase of the cell wall constriction speed with the FtsZ treadmilling speed in *B. subtilis* reflects the lack of a rate-limiting step in regulating the formation of sPG synthase complex, which is consistent with the fact that *B. subtilis* does not have FtsN. We, therefore, suggest that the FtsZ treadmilling-mediated regulation of sPG synthase can adapt its kinetic scheme to meet the distinctive functional needs of cell division in different bacteria.

Our work opens more questions than answers. First, additional rate-limiting factors may render the insensitive dependence of total sPG synthesis activity on FtsZ treadmilling speeds. For instance, the abundance of cell wall-building material, lipid II, may limit cell wall synthesis in *E. coli* (58–61). We will explore this possibility in our future studies. Second, super-fission (SF) variants of FtsA, FtsQLB, or FtsWI can either completely or partially bypass FtsN for cell division; consequently, sPG synthesis can be activated prematurely, resulting in shortened daughter cells in *E. coli* (17, 48, 62). What molecular mechanisms enable these SF mutants to persistently activate sPG synthases and complete the septal constriction without FtsN? It is likely that in these SF mutants, the kinetic rates of sPG synthase activation are altered, which, according to our model (**Figs. 2C and S3**), will change the dependence of sPG synthesis on FtsZ treadmilling speed. Additionally, septum morphology may be altered, as the persistence run length of these variant complexes on the Z-track or the sPG track would differ. Investigating these predictions may shed light on the role(s) of FtsZ treadmilling speed dependence of sPG synthesis in cell division. Third, for simplicity, our model assumes FtsWI/QLB and FtsN to have the same FtsZ-binding potentials. These binding potentials are mediated by the two essential membrane tethers (*i.e.*, FtsA and ZipA) in the Z-ring with distinctive molecular bases: FtsN engages with FtsZ only *via* FtsA (47–50), whereas FtsWI/QLB binds to FtsZ through either FtsA or ZipA (2, 7, 9, 10). Further work is required to investigate how these potentially distinctive FtsZ-binding influence the sPG synthase formation and why the two-tether system exists in the first place. Lastly, in the absence of FtsZ, a sufficient septal level of FtsN is required to sustain the necessary sPG synthesis for the later stage of septum constriction, as predicted by our model. How does the spatial-temporal pattern of FtsN evolve at the septum and affect the sPG synthesis? What controls the septal accumulation of FtsN? It is known that FtsA mediates the septal recruitment of FtsN from both the cytoplasmic and periplasmic sides, the former mediated by direct interaction between FtsN and FtsA (47–50), and the latter by the recruitment of FtsEX and EnvC (2, 7, 9, 10), which leads to amidase activation to create denuded PG for FtsN to bind (22, 25–27). Interestingly, denuded PGs are also degraded by hydrolases (20–22), which offer a mechanism to couple old sPG degradation with sPG synthesis through FtsN (28). How do the amidases and hydrolases synchronize their actions to underlie FtsN’s spatial-temporal distribution pattern? How does the degradation of old sPG coordinate with the synthesis of new sPG to constrict the septum robustly without potential lesions in time and space? Ultimately, all the above questions boil down to the fundamental and unanswered question: What is the coherent mechanism that orchestrates the spatial-temporal coordination between divisome proteins to ensure robust cell division? The two-track model presented in this work will serve as the starting point to allow further explorations of these exciting questions.

## Materials and Methods

Modeling details are presented in the SI.

### Experimental Methods

#### Activation lifetime measurements

To obtain lifetimes of cell wall synthase molecules undergoing activation, we utilized single-molecule tracking data of Halo-FtsI^SW^ in live *E. coli* cells labeled with ZapA-mNeoGreen (Strain JM140) (34) and vertically oriented in nanoholes of M9-agarose gel pad. Segmented trajectories of single Halo-FtsI^SW^ molecules were categorized using 15 nm/s as a cut off, where segments below 15 nm/s were classified as slow moving on the sPG-track, and segments above 15 nm/s were classified as fast moving on the Z-track. Segments that did not show statistically significant processivity were classified as immobile (35). We then pooled segments that were on the Z-track initially and then transitioned to the sPG track or the immobile state to obtain their lifetimes in sec on the Z-track (Fig. 2D). These lifetimes reflect the time it takes for an FtsI molecule to exit the Z-track to become primed (immobile) or active (move on the slow sPG track).

#### Coefficient variation (CV) from HADA labelled cells

To determine CV from HADA labelled cells, we utilized images from (29). Briefly, cells were labelled with HADA for 8 min followed by ethanol fixation and image acquisition (see (29) for details). Fluorescence images of cells were segmented and two-line scans of fluorescence intensity per cell, one along the septa, the other along the quarter position of the cell were measured using ImageJ. The mean background value per pixel calculated from the quarter cell line scan was then subtracted from the fluorescence intensity value at each pixel from the line scan at the septum using a custom MATLAB code. The CV of the background-subtracted septal line scan was determined by calculating the standard deviation of the mean divided by the mean of the line scan.

#### Analysis of FtsW speeds in relation to cell diameter

FtsW-TagRFP velocities were examined in relation to cell diameter based on the data obtained in (35). Briefly, 2D-SMT was performed on WT cells expressing FtsW-TagRFP (BW25113) in M9 agarose. Cell diameter was used as a measure of cell division progression where velocities were binned according to the corresponding diameter of the cells.

##### His_6_-FtsN^SPOR^ purification

His_6_-FtsN^SPOR^ was purified from *E. coli* SHuffle® T7 (New England Biolabs) as described (63). PG sacculi from *Bacillus subtilis* from strain PY79 were purified as described (26).

##### FtsN^SPOR^ labeling by Cy3B

FtsN^SPOR^ labeling was performed by incubating the purified protein with 1:1 ratio of Mono-Reactive NHS Ester Cy3B dye (PA63101, Cytiva) in 100 mM NaHCO3/Na2CO3 buffer (pH 8.3) at 25°C for 2hr. The free dye was removed using a PD-10 desalting column (17085101, Cytiva) with 100 mM PBS (pH 7.0) to elute the labelled protein. The labeling ratio of Cy3B-FtsN^SPOR^ is 0.9 (dye-to-protein).

##### Cy3B-FtsN^SPOR^ binding to sacculi from *B. subtilis*

100 ul of 1:10 diluted sacculi was added into the well of Nunc™ Lab-Tek™ II Chambered Coverglass (155382PK, Thermo Fisher Scientific). Sacculi were allowed to adhere for 10 min; then the suspension was removed by aspiration, and well was washed three times with 200 μL of 1xPBS to remove any nonadherent sacculi. The well was allowed to air dry for 5 min. Then 100 ul of different concentrations of Cy3B-FtsN^SPOR^ solution (0, 0.1nM, 1nM, 3nM, 10nM, 30nM, 100nM, 300nM, 1uM, 3uM, 10uM) were added into the well, and incubate for 10 min. Unbound protein was removed by washing the wells three times with 200 μL of PBS. Finally, 200 μL of PBS was added and sample was examined immediately under the microscope.

## Author Contributions

J.X., D.S.W., and J.L. conceived and designed the study. L.H.H and T.N. performed the computations of the mathematical model. A.P., Z.X.L, and A.Y. performed the experiments. L.H.H., A.P., T.N., Z.X.L, A.Y., D.S.W., J.X., and J.L. analyzed the data, wrote, reviewed, and edited the manuscript.

## Acknowledgements

This work was supported by start-up funds from the Johns Hopkins University School of Medicine, Johns Hopkins Catalyst award, the National Science Foundation (2105837 and 2148534) to J.L., NIGMS R35GM136436 to J.X., NIH R01GM125656 to D.S.W., NIGMS F32GM150262 to A.P., and T32 NIH training grant (GM135131) and NSF graduate research fellowship to T.N. Computing work was carried out at the Advanced Research Computing at Hopkins (ARCH) core facility (rockfish.jhu.edu), which is supported by the National Science Foundation (NSF) grant number OAC1920103. The funders had no role in study design, data collection and analysis, decision to publish, or preparation of the manuscript.

## Figure & Captions

**Figure S1.**
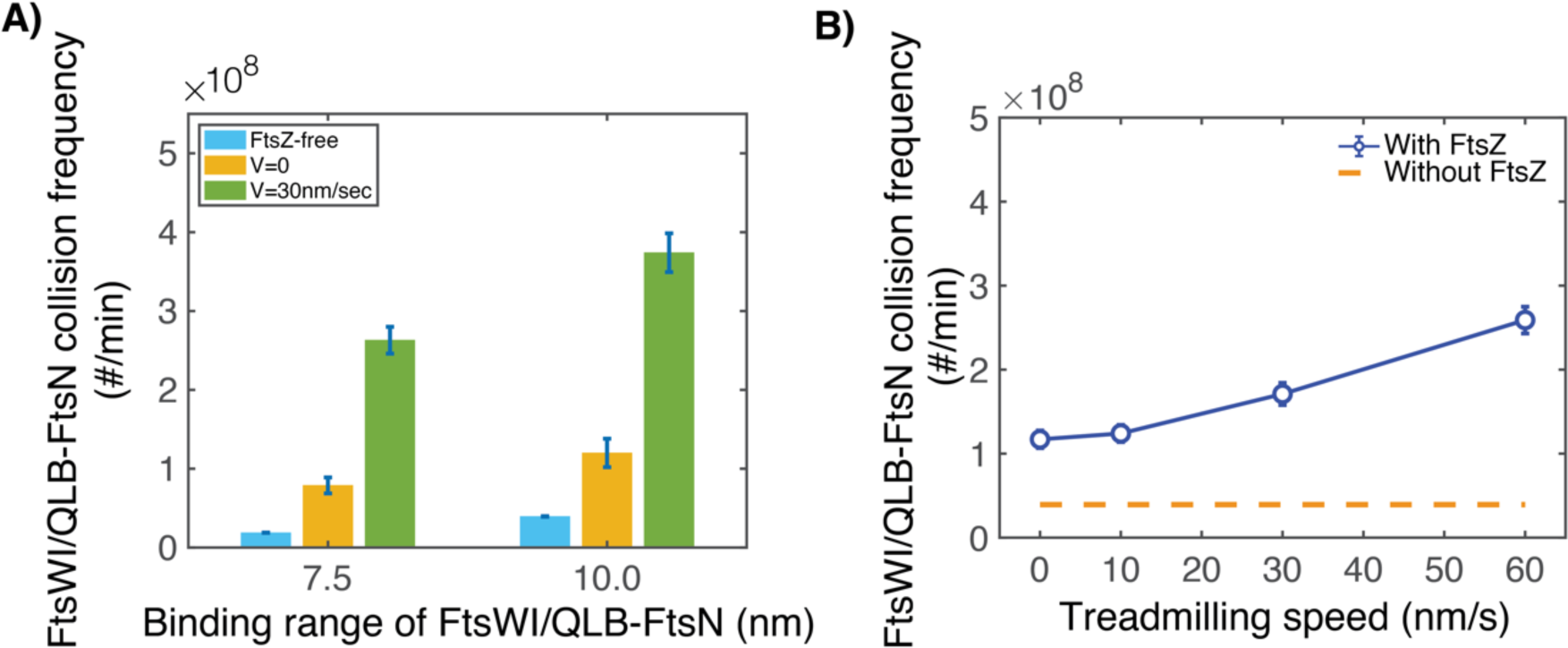
FtsZ treadmilling-mediated enhancement of collision frequency remains robust against the variation of model parameters. A) The enhancement effect holds against the changes in the threshold distance that defines close contact. FtsZ polymer treadmilling speed is fixed at 30 nm/s. B) The enhancement effect holds against the changes in the free diffusion constants of FtsWI/QLB and FtsN. Here, the free diffusion constants of FtsWI/QLB and FtsN are chosen to be 0.1 um^2^/s, as compared to 0.04 um^2^/s as our nominal case in the main text.

**Figure S2.**
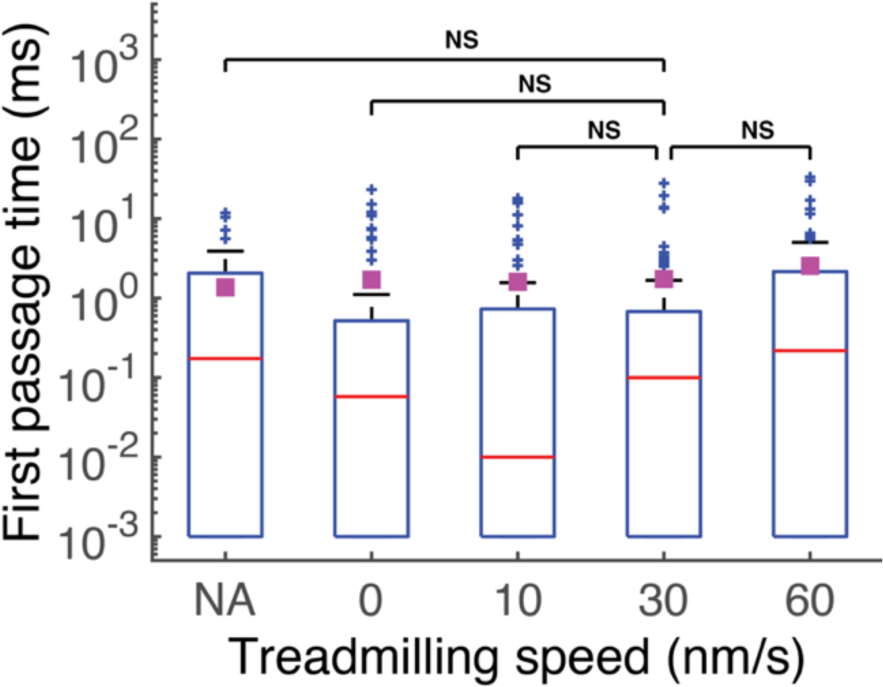
First-passage-time from model calculation shows that FtsWI/QLB and FtsN come into a close contact is insensitive to the presence of FtsZ.

**Figure S3.**
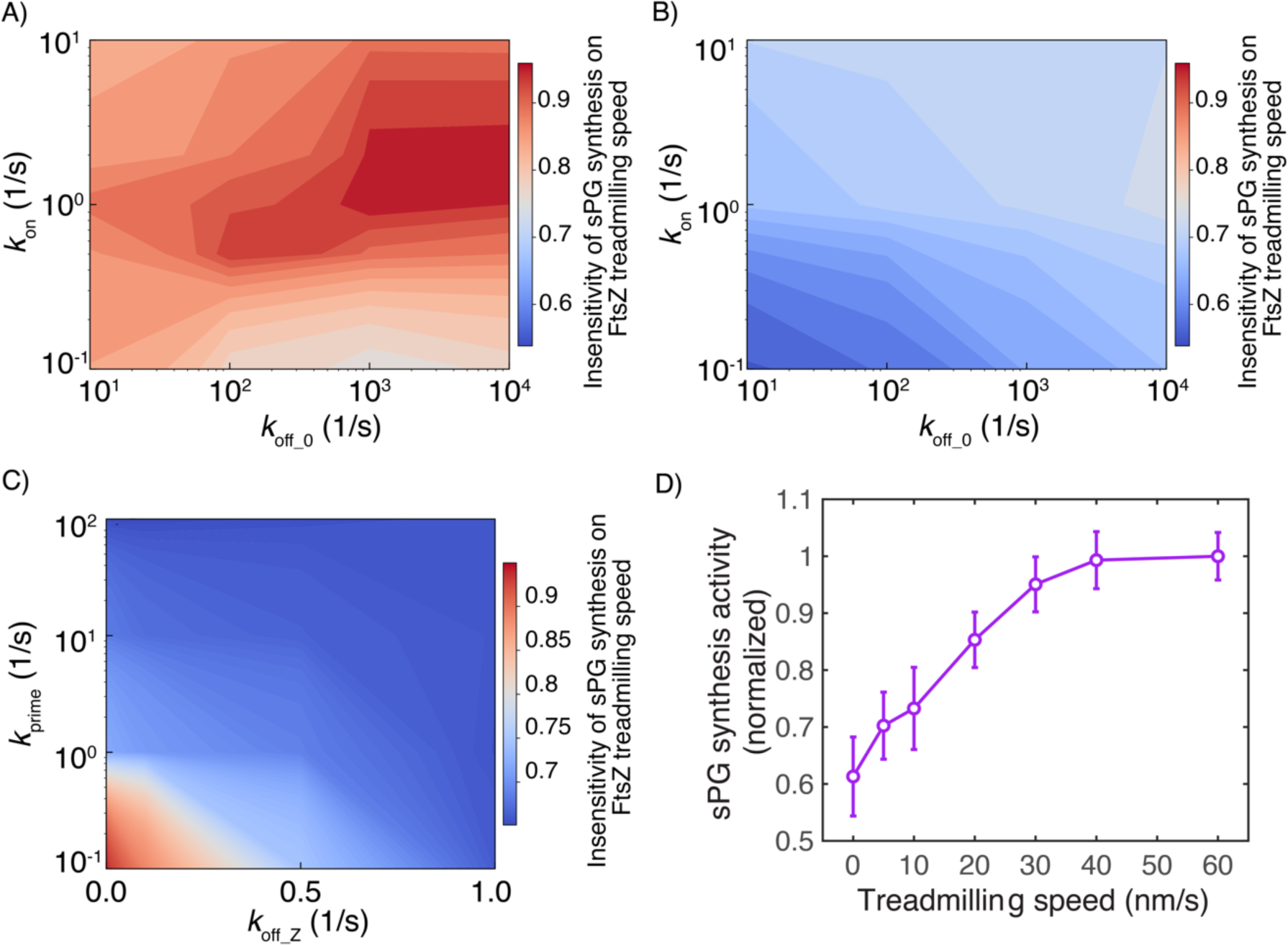
Formation kinetics of sPG synthase complex modulates the dependence of sPG synthesis on FtsZ treadmilling speed. A) Model phase diagram calculation showing that the dependence of sPG synthesis activity on FtsZ treadmilling speed modulated by *k*_on_ and *k*_off_0_ when *k*_prime_ = 0.1/s. B) Model phase diagram calculation showing that the dependence of sPG synthesis activity on FtsZ treadmilling speed modulated by *k*_on_ and *k*_off_0_ when *k*_prime_ = 1/s. C) Model phase diagram calculation showing that the dependence of sPG synthesis activity on FtsZ treadmilling speed modulated by *k*_prime_ and *k*_off_Z_. In A-C), if not otherwise mentioned, all the other model parameters were fixed in accordance with their nominal values listed in the model parameter table. For each parameter pair in the phase diagram calculations, we computed the corresponding curve of normalized sPG synthesis activity vs FtsZ treadmilling speed, from which we determined the sPG synthesis activity at the FtsZ treadmilling speed of 5 nm/s relative to that at 30 nm/s. This way, this normalized sPG synthesis activity defines the value of the colormap. Since the normalized sPG synthesis activity was observed to only drop to ∼ 90% as the FtsZ treadmilling speed decreases from ∼ 30 nm/s to ∼ 5 nm/s in *E. coli*, we used the 90% as the quantitative criterium: if the normalized sPG synthesis activity at the FtsZ treadmilling speed of 5 nm/s is ≥ 90%, then we determine that the sPG synthesis activity insensitively depends on FtsZ treadmilling speed; otherwise, sensitive. Taken the multiple threads together, A-C) show that the only insensitive zone of sPG synthesis against FtsZ treadmilling speed (the dark-red regime) is with an intermediate *k*_on_, a large *k*_off_0_, a small *k*_prime_, and an even smaller *k*_off_Z_. This finding suggests 1) the insensitive dependence of sPG synthesis on FtsZ treadmilling speed entails the rate-limiting step of the forming active sPG synthase on Z-track to be the step of FtsWI/QLB-FtsN^‡^→FtsWI/QLB-FtsN*, and 2) to synthesize enough sPG requires a sufficiently long Z track-bound time of FtsWI/QLB-FtsN^‡^ so that it has the chance of being converted to FtsWI/QLB-FtsN*. D) sPG synthesis activity sensitively depends on FtsZ treadmilling speed when the priming step is not rate-limiting. In this case, *k*_prime_ = 100/s.

**Figure S4.**
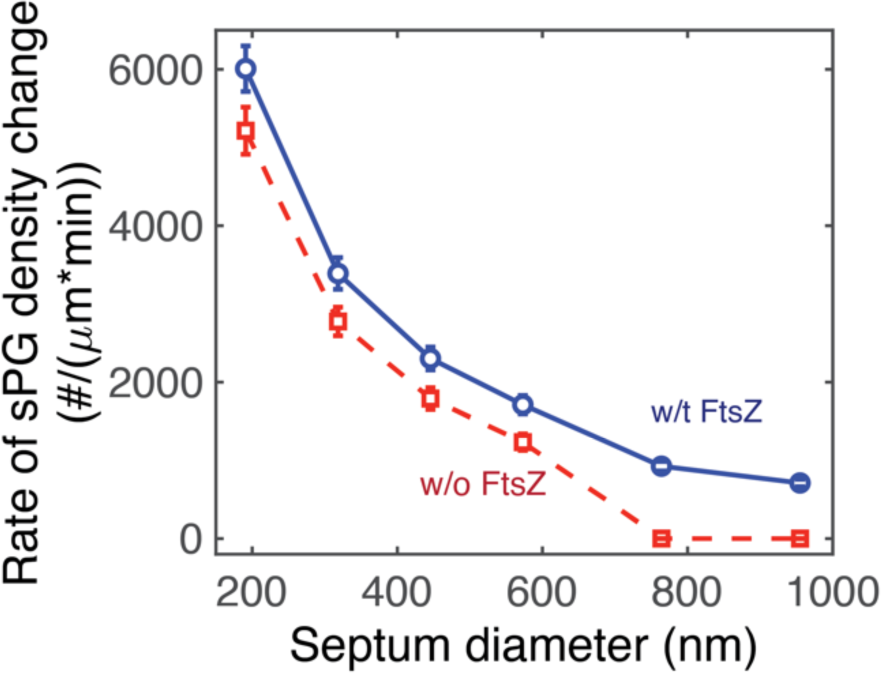
Sustained sPG synthesis and septum constriction by FtsN at the late stages of septum constriction persists as long as FtsN binds to denuded PG tightly. Hereby, *k*_a_, the rate by which FtsN binds to the denuded PG is decreased to 0.2/s from 1.0/s as that in Fig. 4. Despite the decrease in *k*_a_, not only the overall rate of sPG synthesis and septum constriction is unchanged, but also FtsZ remains largely dispensable in the later stages of septum constriction (*i.e.*, the septum diameter < 600 nm).

**Figure S5.**
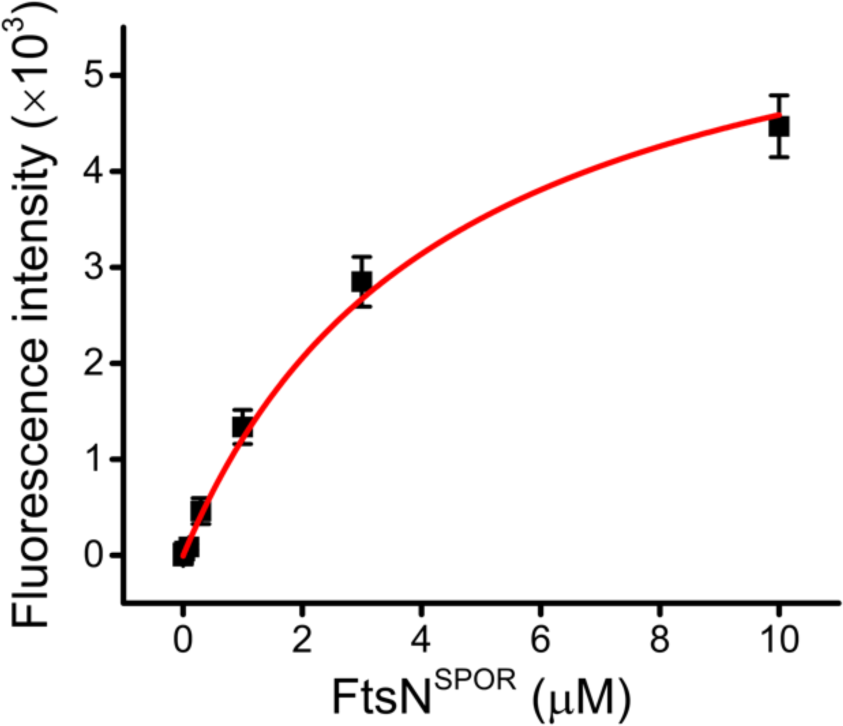
Experimental measurements show strong binding affinity between FtsN and denuded PG binding using purified sacculi. By fitting this data to Hill function, the dissociation constant *K*_D_ for FtsN-denuded PG binding is calculated to be ∼ 4.4 ± 0.9 μM. Please see the Methods and Materials section for the details of this biochemistry experiments.

## Movie Captions

**Movie S1.** Representative movie of 30 seconds of our simulation across the full septum sans reactions. The simulation herein includes 20 FtsWI/QLB complexes and 20 FtsN particles, which are depicted as blue discs and pink discs, respectively. Treadmilling FtsZ filaments are depicted in purple moving at 30 nm/s. Complex coordinates were plotted every 0.1 seconds. The FtsZ filaments and particles are scaled up in size by a factor of two to aid the ease of visualization.

**Movie S2.** Zoom in view of a bundle of FtsZ corralling together an FtsWI/QLB complex (blue disc) and FtsN particle (pink disc) over the course of 20 seconds with the same simulation conditions and visualization properties as Movie 1.

**Movie S3.** Representative movie of sPG synthesis activity from simulating the two-track model in the Stage I. Hereby, the nominal model parameter set is chosen in accordance with the model parameter table (Table S1). Within this 60-second movie, the colormap of the synthesized sPG follows that in the Figure 3A: *i.e.*, lighter colored sPG strands were synthesized earlier in the simulation than the darker colored strands. Upon dissociating from FtsZ polymers, the FtsWI/QLB can be seen staying stationary for some amount of time, which is assumed to attach to the denuded PG, prior to the initiation of sPG synthesis at a velocity of 8 nm/s in the left or right direction. FtsZ is shown to be a lighter purple for clarity.

## Supplemental Information

### Modeling Methods

Our computational model treats the septum as a 2D rectangular region, where the periodic and reflective boundary conditions are applied lengthwise and widthwise, respectively. This simulation domain has different layers (schematized in Figs. 2 and 4), which are occupied by Z-track, periplasm, and sPG-track separately so that they do not physically overlap.

#### On Z-track

There are many FtsZ filaments. All the FtsZ filaments are positioned parallel to the x-direction (lengthwise). To model the FtsZ clusters observed in the Z-ring *in vivo* (23, 28), we grouped the FtsZ filaments into the bundles composed of multiple FtsZ filaments positioned side by side along the y-direction (widthwise). FtsZ filaments undergo the treadmilling process, which is modeled deterministically, similar to the previous work (34): an FtsZ filament grows by adding one subunit to its growing end at a time, with the time interval being determined by the size of the FtsZ subunit and the treadmilling speed; the subunit on the shrinking end of the FtsZ filament is removed at the same rate as the growing end to keep the length of the FtsZ filament fixed.

#### In periplasm

There are FtsWI/QLB and FtsN molecules that can form FtsWI/QLB-FtsN complex. These molecules are modeled as spherical particles that undergo the reaction-diffusion processes and switch between the states that is bound to Z-track, sPG-track, or freely diffuse in periplasm. The reactions shown in the schematic reaction pathway of the model (Figs. 2 and 4) are simulated as Poisson process, using the kinetic Monte Carlo approach. The reaction rates used in the simulation are listed in Table S1.

We used Brownian dynamics formulism to simulate the movement of the particles, which is governed by the overdamped Langevin equation: ξ*d*r/dt = *f*(t)+*ρι*(t). Here, *r* is the position of the particle, ξ is the friction coefficient, *f* is the net external forces that stems from the binding potentials and excluded volume repulsion (see below), and η represents the random force resulting from the thermal motion of the solvent molecules. The random force, η satisfies 〈η_*i*_(*t*)〉 = 0 and 〈η_*i*_(*t*)η_#_(*t*^$^)〉 = 2ξ*k*_*B*_Tδ_*ik*_δ(*t* − *t*^$^), where δ_*i*#_ is the Kronecker delta and δ(*t* − *t*^$^) is the Dirac delta function. Similar to the previous work (34), we used the short-ranged harmonic potential to model the binding potentials between FtsZ and FtsWI/QLB, FtsN, and FtsWI/QLB-FtsN, respectively, which are characterized by the binding energy and the binding range (see Table S1 below). To prevent the overlap of the particles, the excluded volume repulsion between the particles is applied. The simulation time step was set to be 1 α*s*.

#### On sPG track

One active sPG synthase produces one sPG strand at a time at the rate of 8 nm/s. The direction of individual sPG synthesis event has the equal probability of being left or right; but once the synthesis event starts, its direction remains fixed until the end. For the sake of simplicity, the sPG strand is assumed to be parallel to the x-axis without explicating the crosslinking events. Concurrent with the sPG synthesis event, the active sPG synthase is assumed to move directionally at the tip of synthesizing sPG strand with the same velocity of 8 nm/s.

---

**Table S1.**
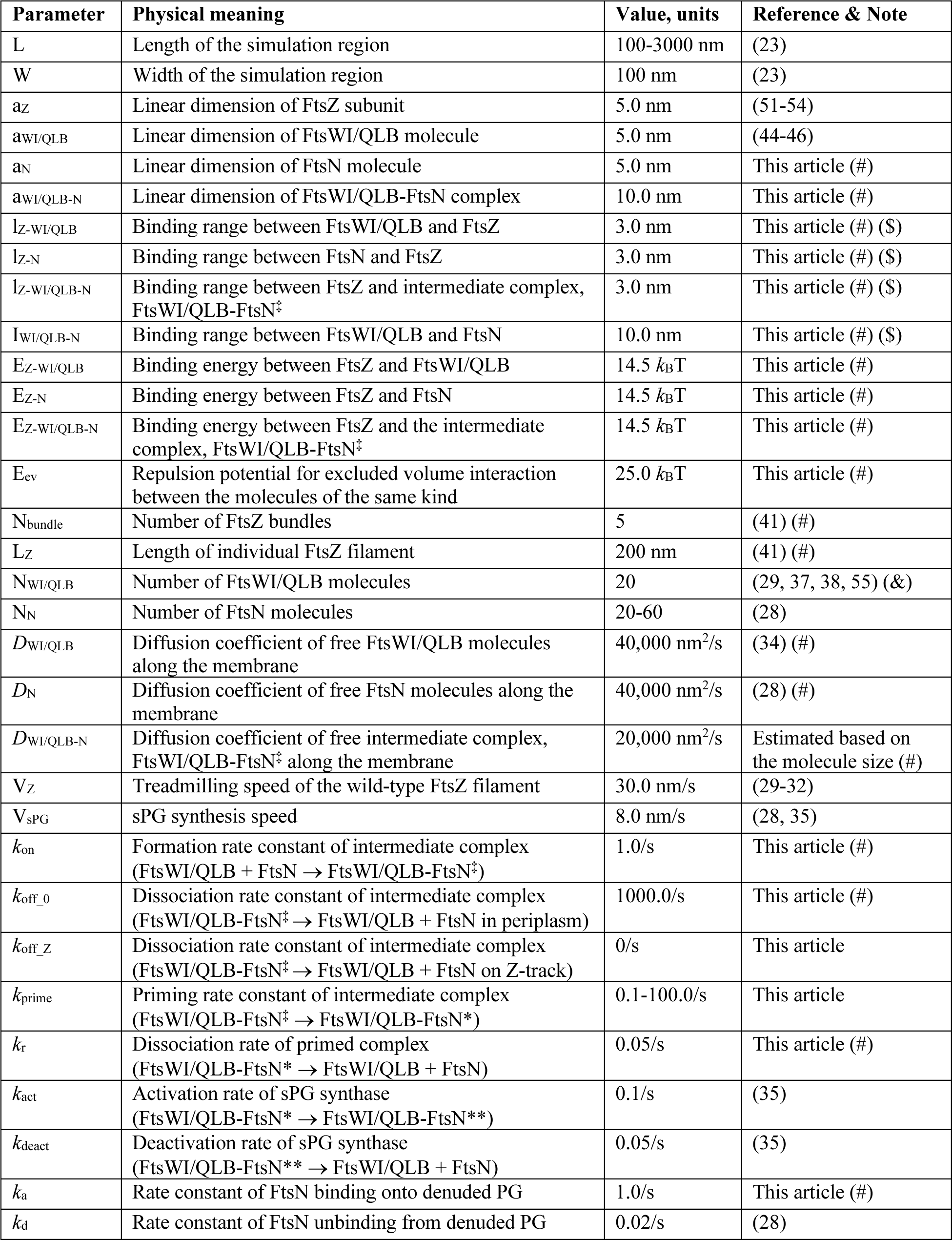
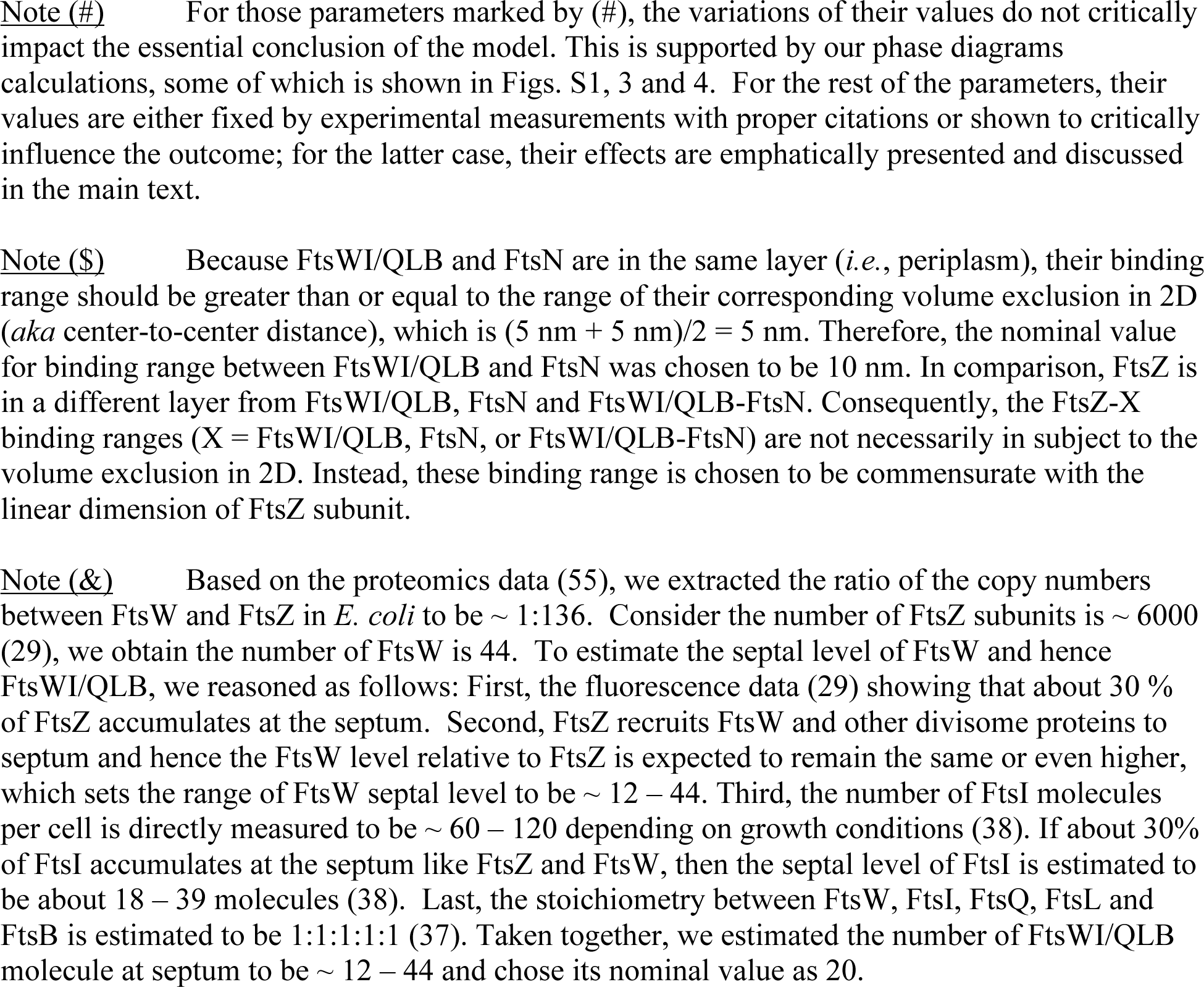
Model Parameters

| Parameter | Physical meaning | Value, units | Reference & Note |
| --- | --- | --- | --- |
| L | Length of the simulation region | 100-3000 nm | (23) |
| W | Width of the simulation region | 100 nm | (23) |
| az | Linear dimension of FtsZ subunit | 5.0 nm | (51-54) |
| a<sub>WI/QLB</sub> | Linear dimension of FtsWI/QLB molecule | 5.0 nm | (44-46) |
| a<sub>N</sub> | Linear dimension of FtsN molecule | 5.0 nm | This article (#) |
| a<sub>WI/QLB-N</sub> | Linear dimension of FtsWI/QLB-FtsN complex | 10.0 nm | This article (#) |
| l<sub>Z-WI/QLB</sub> | Binding range between FtsWI/QLB and FtsZ | 3.0 nm | This article (#) (\$) |
| l<sub>Z-N</sub> | Binding range between FtsN and FtsZ | 3.0 nm | This article (#) (\$) |
| l<sub>Z-WI/QLB-N</sub> | Binding range between FtsZ and intermediate complex, FtsWI/QLB-FtsN<sup>‡</sup> | 3.0 nm | This article (#) (\$) |
| l<sub>WI/QLB-N</sub> | Binding range between FtsWI/QLB and FtsN | 10.0 nm | This article (#) (\$) |
| E<sub>Z-WI/QLB</sub> | Binding energy between FtsZ and FtsWI/QLB | 14.5 k<sub>B</sub>T | This article (#) |
| E<sub>Z-N</sub> | Binding energy between FtsZ and FtsN | 14.5 k<sub>B</sub>T | This article (#) |
| E<sub>Z-WI/QLB-N</sub> | Binding energy between FtsZ and the intermediate complex, FtsWI/QLB-FtsN<sup>‡</sup> | 14.5 k<sub>B</sub>T | This article (#) |
| E<sub>ev</sub> | Repulsion potential for excluded volume interaction between the molecules of the same kind | 25.0 k<sub>B</sub>T | This article (#) |
| N<sub>bundle</sub> | Number of FtsZ bundles | 5 | (41) (#) |
| L<sub>Z</sub> | Length of individual FtsZ filament | 200 nm | (41) (#) |
| N<sub>WI/QLB</sub> | Number of FtsWI/QLB molecules | 20 | (29, 37, 38, 55) (&) |
| N<sub>N</sub> | Number of FtsN molecules | 20-60 | (28) |
| D<sub>WI/QLB</sub> | Diffusion coefficient of free FtsWI/QLB molecules along the membrane | 40,000 nm<sup>2</sup>/s | (34) (#) |
| D<sub>N</sub> | Diffusion coefficient of free FtsN molecules along the membrane | 40,000 nm<sup>2</sup>/s | (28) (#) |
| D<sub>WI/QLB-N</sub> | Diffusion coefficient of free intermediate complex, FtsWI/QLB-FtsN<sup>‡</sup> along the membrane | 20,000 nm<sup>2</sup>/s | Estimated based on the molecule size (#) |
| V<sub>Z</sub> | Treadmilling speed of the wild-type FtsZ filament | 30.0 nm/s | (29-32) |
| V<sub>sPG</sub> | sPG synthesis speed | 8.0 nm/s | (28, 35) |
| k<sub>on</sub> | Formation rate constant of intermediate complex (FtsWI/QLB + FtsN → FtsWI/QLB-FtsN<sup>‡</sup>) | 1.0/s | This article (#) |
| k<sub>off_0</sub> | Dissociation rate constant of intermediate complex (FtsWI/QLB-FtsN<sup>‡</sup> → FtsWI/QLB + FtsN in periplasm) | 1000.0/s | This article (#) |
| k<sub>off_Z</sub> | Dissociation rate constant of intermediate complex (FtsWI/QLB-FtsN<sup>‡</sup> → FtsWI/QLB + FtsN on Z-track) | 0/s | This article |
| k<sub>prime</sub> | Priming rate constant of intermediate complex (FtsWI/QLB-FtsN<sup>‡</sup> → FtsWI/QLB-FtsN*) | 0.1-100.0/s | This article |
| k<sub>r</sub> | Dissociation rate of primed complex (FtsWI/QLB-FtsN* → FtsWI/QLB + FtsN) | 0.05/s | This article (#) |
| k<sub>act</sub> | Activation rate of sPG synthase (FtsWI/QLB-FtsN* → FtsWI/QLB-FtsN**) | 0.1/s | (35) |
| k<sub>deact</sub> | Deactivation rate of sPG synthase (FtsWI/QLB-FtsN** → FtsWI/QLB + FtsN) | 0.05/s | (35) |
| k<sub>a</sub> | Rate constant of FtsN binding onto denuded PG | 1.0/s | This article (#) |
| k<sub>d</sub> | Rate constant of FtsN unbinding from denuded PG | 0.02/s | (28) |

Note (#) For those parameters marked by (#), the variations of their values do not critically impact the essential conclusion of the model. This is supported by our phase diagrams calculations, some of which is shown in Figs. S1, 3 and 4. For the rest of the parameters, their values are either fixed by experimental measurements with proper citations or shown to critically influence the outcome; for the latter case, their effects are emphatically presented and discussed in the main text.
Note (\$) Because FtsWI/QLB and FtsN are in the same layer (*i.e.*, periplasm), their binding range should be greater than or equal to the range of their corresponding volume exclusion in 2D (*aka* center-to-center distance), which is $(5\text{ nm} + 5\text{ nm})/2 = 5\text{ nm}$ . Therefore, the nominal value for binding range between FtsWI/QLB and FtsN was chosen to be 10 nm. In comparison, FtsZ is in a different layer from FtsWI/QLB, FtsN and FtsWI/QLB-FtsN. Consequently, the FtsZ-X binding ranges (X = FtsWI/QLB, FtsN, or FtsWI/QLB-FtsN) are not necessarily in subject to the volume exclusion in 2D. Instead, these binding range is chosen to be commensurate with the linear dimension of FtsZ subunit.
Note (&) Based on the proteomics data (55), we extracted the ratio of the copy numbers between FtsW and FtsZ in *E. coli* to be $\sim 1:136$ . Consider the number of FtsZ subunits is $\sim 6000$ (29), we obtain the number of FtsW is 44. To estimate the septal level of FtsW and hence FtsWI/QLB, we reasoned as follows: First, the fluorescence data (29) showing that about 30 % of FtsZ accumulates at the septum. Second, FtsZ recruits FtsW and other divisome proteins to septum and hence the FtsW level relative to FtsZ is expected to remain the same or even higher, which sets the range of FtsW septal level to be $\sim 12 - 44$ . Third, the number of FtsI molecules per cell is directly measured to be $\sim 60 - 120$ depending on growth conditions (38). If about 30% of FtsI accumulates at the septum like FtsZ and FtsW, then the septal level of FtsI is estimated to be about $18 - 39$ molecules (38). Last, the stoichiometry between FtsW, FtsI, FtsQ, FtsL and FtsB is estimated to be 1:1:1:1:1 (37). Taken together, we estimated the number of FtsWI/QLB molecule at septum to be $\sim 12 - 44$ and chose its nominal value as 20.

## Notes

### Competing Interest Statement

The authors have declared no competing interest.

